# Time Series GWAS for Iron Deficiency Chlorosis Tolerance in Soybean using Aerial Imagery

**DOI:** 10.1101/2025.05.30.657051

**Authors:** Matthew E. Carroll, Ashlyn Rairdin, Liza Van der Laan, Antonella Ferela, Sahishnu Hanumolu, Soumik Sarkar, Baskar Ganapathysubramanian, Arti Singh, Asheesh K. Singh

## Abstract

The use of drones has become a commonly used tool by plant scientists to aid in plant phenotyping endeavors. Iron deficiency chlorosis (IDC) is a commonly observed abiotic stress in soybean fields with high soil pH levels. IDC severity is visually classified, and recent work has shown that digital imaging techniques using both ground and UAS-acquired imagery can be utilized for automated severity ratings. In our study, we compared the classification accuracy of two flight altitudes to determine the optimal flight parameters for IDC phenotyping. In addition to this, we investigated the ability to use image-predicted scores for genome wide association study (GWAS), as well as the effect of IDC on traits such as canopy area and canopy growth and development. We also report a tool for semi-automated plot extraction from orthomosaic images that can be easily integrated with UAS. We noted that 43 days after planting was an ideal time for IDC severity ratings as the highest number of significant SNPs were reported at this timepoint.

## 1 Introduction

Recent advances in stress phenotyping in crop species include sensors carried on drones (Herr et al. 2023; Asheesh K Singh et al. 2021); with a well-established pipeline containing multiple steps from data collection, data transfer, cyberinfrastructure, data processing, data analytics, and interpretation followed by actionable outcomes (Guo et al. 2021). Advances in phenotyping capabilities that integrate machine learning (ML) and object detection can significantly reduce the cost and time to phenotype plant traits (**riera2021deepz**; Akintayo et al. 2018; Chiranjeevi, Sadaati, et al. 2023; Gao et al. 2018; Ghosal, Zheng, et al. 2019; Kar, Nagasubramanian, Elango, Carroll, et al. 2023; Saadati et al. 2023). It also enables a time-series investigation of plant traits and their response to stressors. Furthermore, advances in sensors have ushered in deeper insights nto trait features, including for wavelengths outside of the spectrum that the human eye detects. For example, researchers have detected disease signatures of charcoal rot (*Macrophomina phaseolina*) in soybean [*Glycine max* (L.) Merr.] using hyperspectral imaging (Nagasubramanian, S. Jones, Sarkar, et al. 2018), where they deployed a novel 3D deep convolutional neural network (DCNN) assimilating hyperspectral data and interrogated the model to produce physiologically meaningful explanations (Nagasubramanian, S. Jones, Asheesh K Singh, et al. 2019). For crop yield in conjunction with crop health traits, researchers have utilized various sensors in a time series to obtain trait prediction that allows the selection of superior varieties (Chattopadhyay et al. 2023; Aditya Gupta and Asheesh Singh 2023; K. Parmley et al. 2019). Although there are numerous applications of drone-based imaging and with a multitude of sensors, one of the more rapid application areas is in the phenotyping of crop stress, particularly in conjunction with ML models (Arti Singh, S. Jones, et al. 2021).

In the context of plant stress phenotyping, ML and deep learning (DL) can be deployed on four broad categories of problems: Identification, Classification, Quantification, and Prediction (ICQP) (Arti Singh, Ganapathysubramanian, et al. 2016; Asheesh Kumar Singh et al. 2018). In the first stage, researchers may have an interest in simply identifying the stress, followed by a category rating, i.e., classification. These first two stages of the ICQP paradigm are helpful for plant breeding and scouting applications, and more broadly, phenotyping of stress-related traits that are critical in crop breeding (Asheesh K Singh et al. 2021; D. P. Singh et al. 2021). One of the significant challenges of traditional phenotyping methods is the reliance on human raters, which can be mired with subjectivity and suffers from scalability. The inconsistency in manual ratings can have an effect on large scale meta analysis of multiple diseae and stress ratings (J. M. Shook et al. 2021). Since plant breeding programs routinely assess hundreds to thousands of accessions for stress traits, and crop scouts must scan large tracts of land, high throughput phenotyping (HTP) has become necessary (Lobos et al. 2017; Reynolds et al. 2020; Asheesh K Singh et al. 2021; Yang et al. 2017). One of the most versatile tools for HTP is drone-based imagery that is applied to plant stress phenotyping (Duddu et al. 2019; Hatton et al. 2019; Romero et al. 2018). There are multiple uses of drone-based red, green, and blue (RGB) image phenotyping in soybean, for example, foliar disease detection (Castelão Tetila et al. 2017), abiotic stress detection (Dobbels and Lorenz 2019), height (Maimaitijiang et al. 2019), and maturity (Narayanan et al. 2019; Trevisan et al. 2020). The focus on stress phenotyping will remain important because these stressors can cause substantial crop yield and quality loss (Bradley et al. 2021; D. S. Mueller et al. 2020; Arti Singh, S. Jones, et al. 2021; R. P. Singh et al. 2016). These stresses can be caused by biotic and abiotic factors. Among abiotic stresses, nutrient stress is a major category, which is important in breeding and crop production (D. P. Singh et al. 2021) and maintaining genetic gain (Krause et al. 2023).

Iron Deficiency Chlorosis (IDC) in soybean is an important plant stress and is estimated to cause yearly losses of $260 million (Peiffer et al. 2012). IDC is caused by calcareous soils or high soil pH, which limit iron availability, leading to symptoms of chlorosis. The most cost-effective way to prevent plant damage due to IDC is to breed for tolerance. For this strategy, researchers and breeders need to identify sources of IDC tolerance and understand the genetics of resistance to develop IDC-tolerant varieties. The major source of IDC tolerance has been reported on chromosome 3 of soybean, and this large-effect quantitative trait loci (QTL) controls up to 70% of the genetic variation (Lin et al. 1997). Several genome wide association studies (GWAS) have been utilized to study the genetics of IDC tolerance in soybean using field (Mamidi et al. 2014; Merry et al. 2019; Jiaoping Zhang, Naik, et al. 2017) and greenhouse (Assefa et al. 2020) studies, aiming to identify major and minor effect QTL controlling IDC expression. In addition to visual rating of IDC stress in plants, researchers have used HTP and ML for evaluation and classification of the stress (Dobbels and Lorenz 2019; Hassanijalilian, Igathinathane, Bajwa, et al. 2020; Hassanijalilian, Igathinathane, Day, et al. 2023; J. Li et al. 2020; Naik et al. 2017; Jiaoping Zhang, Naik, et al. 2017). However, there are two main gaps in these studies: (a) drone-based imagery for GWAS to determine if digital imagery can elicit additional information compared to visual ratings, and (b) a time series investigation of IDC stress in conjunction with GWAS to study the variation in genetic control at varying time points. The objectives of this paper were to: (1) compare the effect of flight altitude on the accuracy of classification, (2) study the effect of IDC Ratings on canopy size and growth, and (3) conduct GWAS on multiple time points to determine temporal variation in identified significant loci.

## 2 Materials and Methods

### 2.1 Field Setup and Design

Experiments were conducted in 2018 and 2019, at Iowa State University’s Bruner farm (Boone County, Iowa). In 2018, drone imagery was conducted over two fields, while in 2019, a single field was phenotyped. During the 2018 experiment, the first field (hereon referred as IDC field) is historically an IDC stress screening site due to the consistent expression of IDC each year. The second field historically has never shown IDC stress symptoms, and hereon is referred to as the non-IDC field. The IDC and non-IDC field were located on the same ISU research farm (250 meters apart in distance), providing an opportunity to compare the effect of canopy size and growth on IDC ratings as they are geographically next to each other on the same farm, have contrasting IDC expressions on soybean plants, and no confounding weather parameter differences. The IDC field center in 2018 was located at 93.73469°W 42.00998°N; the non-IDC field was located at 93.73405°W 42.01218°N. The 2018 IDC experiment was grown in a randomized complete block design with five replications. Each block had 731 hill plots planted, for a total of 3,655 plots. Each replication consisted of 723 unique accessions belonging to the soybean mini-core collection in maturity groups 0, I, II, III, and IV (Oliveira et al. 2010), along with checks iron-efficient Clark (PI 584533; IDC tolerance) and iron inefficient IsoClark (PI 547430; IDC susceptible) (Assefa et al. 2020). Four replications of each check were included per block, for a total of 40 check plots throughout the field. In 2018, the non-IDC experiment was planted in a randomized complete block design with six replications. Each block had 506 unique accessions belonging to the soybean mini-core collection in maturity groups 0, I, II, and III (Oliveira et al. 2010), and the checks Clark and Isoclark were also included and replicated three times in each block, for a total of 3,072 plots. In 2018, the IDC and non-IDC field had 477 soybean accessions in common from the soybean mini-core diversity panel (Coser et al. 2017; de Azevedo Peixoto et al. 2017; Moellers et al. 2017). Both IDC and non-IDC experiment plots were planted in a hill plot, with five seeds planted per plot, 3.81 cm spacing between each seed, and 76 cm distance (in each of the four directions - rows and columns) plot to plot distance. The xperiment was hand-planted, with plots arranged in a grid manner. The non-IDC field planting date was May 24th and the IDC field planting date was May 29th. Weeds were controlled manually throughout the growing season, to remove or minimize the confounding effects of weeds on soybean growth and development.

In 2019, the IDC experiment was grown at 93.7339°W 42.0101°N. This field was adjacent to the 2018 IDC field experiment and was planted in a randomized block design with 580 accessions in four replications. Smaller number of accessions were planted due to seed source issues from the prior year, with lines that had less than three plots that germinated in 2018 removed from the 2019 experiment. All 580 accessions grown in 2019 were part of the 2018 field experiment. Clark and IsoClark checks were each planted in four plots per replication, giving a total of 588 plots per replication and a field total of 2,352 plots. Experiment details were the same as previously explained for the 2018 IDC field, with a planting date of June 6th. The non-IDC field was not utilized in 2019.

### 2.2 Manual Ratings

Trained raters conducted manual ratings of the IDC severity of each plot using rating scales previously described in the literature (Rodriguez de Cianzio et al. 1979; Jiaoping Zhang, Naik, et al. 2017). Briefly, a score of 1 showed no symptoms of IDC, and a score of 5 showed severe yellowing, necrotic tissue, and stunting of growth. These ratings served as the ground truth for image-classified scores. Ratings were only taken on the IDC stressed fields and were taken on 31, 40, 43, 54, and 70 (DAP) in 2018, and 21, 33, 43, 47, and 55 DAP in 2019. No IDC stress symptoms were noted in the non-IDC experiment, therefore, no ratings were taken. Five data collection dates were recorded for each season. One time point from each year was removed due to the inconsistent time intervals of data collection. In 2018, 40 days after planting (DAP) was removed, while in 2019, 47 DAP was removed from all analyses.

### 2.3 Drone Flights and Imaging

Aerial phenotyping was conducted using a Matrice 600 pro and the Zenmuse X5 camera with the Olympus 45 mm focal length lens (DJI Technology Co., Shenzhen, China). Flights were flown with an 80% front overlap, and a 70% side overlap. Flights were conducted at 60 and 30 meters, with the ground sampling distance (GSD) for each flight being 0.50 and 0.25 cm/pixels, respectively. Flights for IDC ratings were conducted on the same dates as the manual ratings, between 10 am and 2 pm. Flights for the non-IDC field were taken on 37, 44, 49, 63, and 71 DAP. The non-IDC field imaging data were used for canopy growth comparisons.

### 2.4 Image Processing

#### 2.4.1 Image Pre-processing

Images were stitched using Pix4D (Pix4D, SA). Upon ortho-mosaicing of the field experiment image, a semi-automated pipeline was developed that facilitated plot delineation and canopy area extraction. Briefly, using ArcGIS we obtained the coordinates of the four corners in the field. This was inputted into a CSV file, which was called from the custom Python script for plot extraction. The script then calculated the angle and distance between each corner. Subsequently, in ArcGIS using the arcpy Python library, the script was used to delineate the boundaries of each plot, and labeled with the row-column information for each plot. After plot identification using the automated script described above, three vegetation indices were calculated: Normalized Blue, Green leaf index, and Color index of vegetation (Maimaitijiang et al. 2019). These indices were then used as input for the ISO-clustering classification algorithm. User input was provided to label classes using the ISO-clustering tool. The output from this is the classification of pixels as either plant or background, creating a binary classification. After plant pixels were labeled, each plot was retained with the associated largest connected component that was built based on the labeling done in the earlier step. The largest connected component was then used as a mask to extract canopy information for each plot. The masked RGB image was then converted to the Hue Saturation Value (HSV) color space (*Color Model Conversion function* n.d.). Finally, the color wheel was used to extract Green, Yellow, and Brown pixels from each plot as well as the median Hue value and the canopy area. This data was then exported in a CSV file format for further analysis. Fig. 1 gives an overview of the plot extraction, and image analysis pipeline.

**Figure 1:**
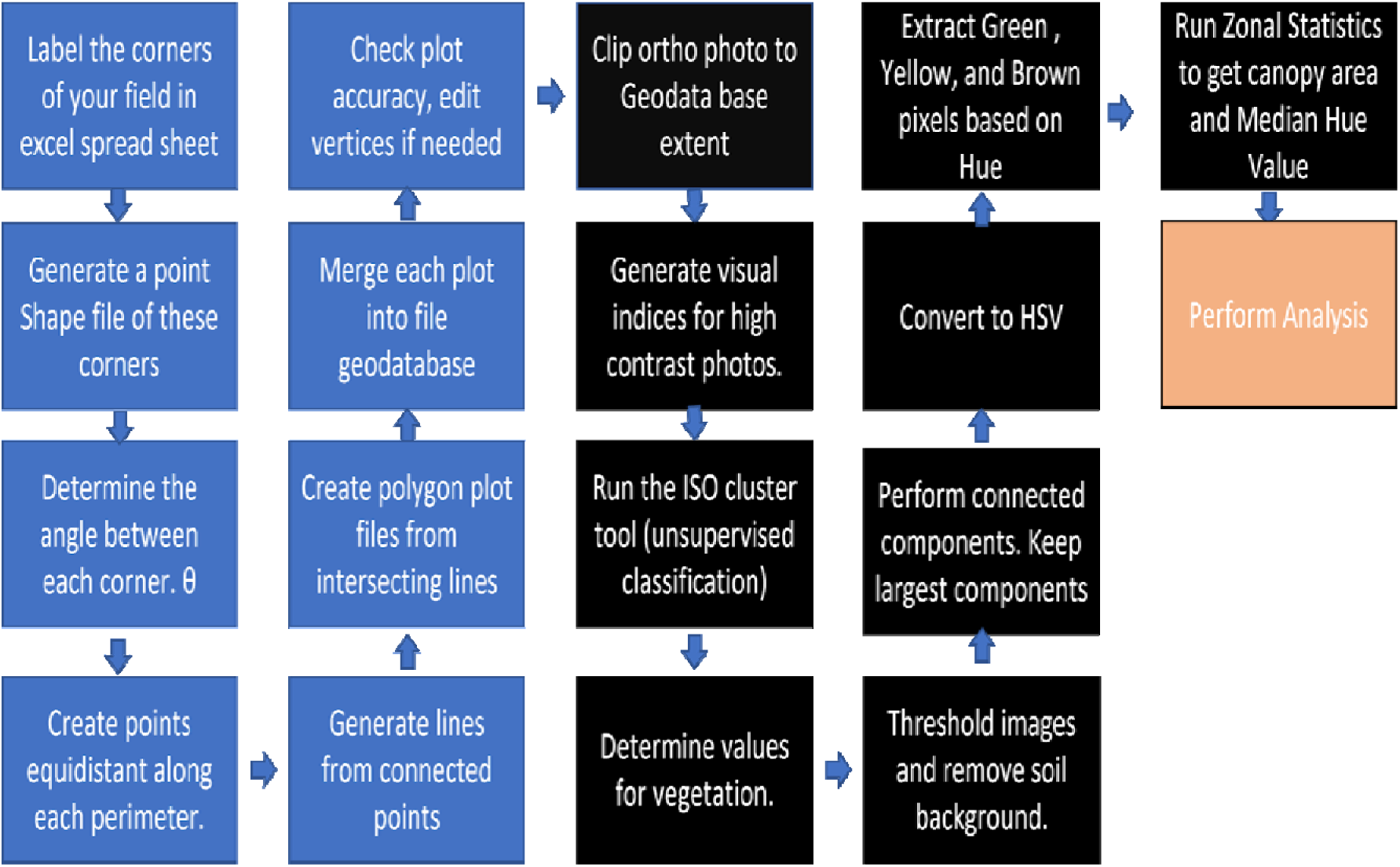
An overview of the image preprocessing steps used to extract data from each plot.

#### 2.4.2 Image Processing - IDC Score Classification

Models were trained using the xgboost package in R to run an extreme gradient boosting machine (Chen and Guestrin 2016) for IDC score classification. The model used the percent green, percent yellow, percent brown, and the median hue value of each canopy to classify each plot. The manual ratings were used as the ground truth to evaluate the accuracy of the models. Accuracy was calculated by taking the sum of the diagonal of the confusion matrix and dividing it by the total number of observations within the testing set. Data sets were subset by height, and flight date. IDC scores for each date were unbalanced, giving a higher likelihood of misclassification of minority IDC classes. To address this issue and increase accuracy for all classifications, weights for each plot were assigned based on the percent of each rating class for a given time point. Each dataset used five-fold cross validation to estimate model accuracy. Five-fold cross validation was repeated six times for each dataset, giving a total of 30 model accuracies using XGBoost. Model tuning parameters used a maximum depth of 15, a learning rate of 0.1, with a maximum of 100 rounds of boosting iterations. The model had a softmax objective funtion for multiclass classification. To determine if there was a statistically significant difference in the accuracy at each timepoint between the 30 and 60 meter flight altitudes, two-sided t-tests were run contrasting the mean accuracy between each flight altitude.

#### 2.4.3 Image Processing - Canopy Traits

Canopy area is presented on a log transformed scale because the canopy growth followed an exponential growth pattern and earlier time point canopy areas were extremely small, making visual comparison difficult. While the time point across the IDC and non-IDC fields (see M and S section) did not coincide at each instance, the time series approach enabled the comparison of canopy growth between IDC and non-IDC fields, and also across years among the IDC fields. In addition to the IDC score severities, three canopy traits were calculated: Canopy Area (CA), Canopy Growth Rate (CGR), and Canopy Reduction (CR). CA was calculated on a per plot basis using equation 1:

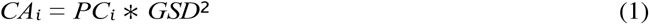

Where *CAi* and *PCi* are the canopy area and pixel count of the the i*th* plot, respectively. GSD is the ground sampling distance. CGR was calculated using equation 2:

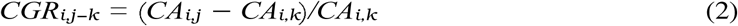

Where *CGR_i,j−k_* is the canopy growth rate of plot i at the *j* − *k* time point, *CA_i,j_* is the canopy area of plot i at the j time-point, and *CAi,k* is the canopy area of plot i at time point k. CGR ordinal comparisons were coded as *T* 1 *T* 2*, T* 2 *T* 3*, T* 3 *T* 4 indicating comparisons of time point1 and time point2, time point2 and time point3, and time point3 and time point4.

CR was only calculated for the 2018 season as there was not a non-IDC field in the 2019 growing season. CR was calculated using the equation 3:

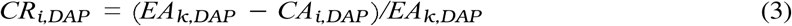

Where *CRi,DAP* is the canopy reduction of plot i on a given DAP, *EAk,DAP* is the expected area of genotype k given DAP, and *CAi,DAP* is the canopy area of plot i at a given DAP. *EAk,DAP* was derived from the data collected from the non-IDC field using the equations 4 and 5:

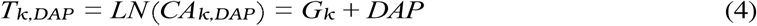

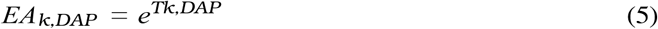

Where *Tk,DAP* is the predicted value of the natural logarithm of the canopy area at a specific DAP, *Gk* is the fixed effect of genotype k, and DAP is days after planting.

### 2.5 GWAS

A Genome Wide Association Study (GWAS) was run on manual IDC ratings, UAV classified IDC ratings, CA, CGR, and CR for all time points. For the IDC severity ratings the response was modeled using an ordinal logistic regression to generate trait best linear unbiased predictors (BLUPs) as described in previous papers with ordinal responses (Moellers et al. 2017). The R package ordinal (Christensen 2022) was used to generate a mixed model treating the genotype as random. IDC severity scores were modeled using equation 6:

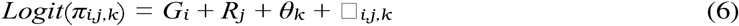

Where π*i,j,k* is the probability of genotype i in replication j to be rated at or below category k, *Gi* is the effect of genotype i, *Rj* is the effect of replication j, and θ*k* is the intercept for severity rating k. [*i,j,k* is the residual for genotype i, in rep j, with a severity rating of k. CA, CGR, and CR BLUPs were calculated using a mixed linear model approach that has been used in previous stress phenotyping research (Coser et al. 2017; Jiaoping Zhang, Arti Singh, et al. 2015) using the lme4 package in R using equation 7:

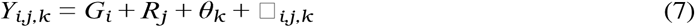

Where *Yi,j,k* is the response of genotype i in replication j with a severity rating of k. *Gi Ri* and θ*k* are the same as described above. Once genotypic BLUPs were calculated for each trait they were used as the phenotypic values for GWAS in Tassel software (Bradbury et al. 2007). SoySNP50K BeadChip was used for the genotypic data (Song et al. 2013), and was downloaded from SoyBase (https://soybase.org). Sites with a minor allele frequency of less than 0.01 were removed, and missing data was imputed using LD KNNi Imputation (Money et al. 2015). A kinship matrix was calculated using Tassel software (Bradbury et al. 2007) using the centered IBS method. PCA was calculated using Tassel and the first three principal components were used to account for population structure in the mixed linear model. A False Discovery Rate (FDR) correction was used with a threshold of p = 0.05 to deem SNPs as significant.

## 3 Results

### 3.1 Drone-based imagery at 30 and 60 Meter flight heights

IDC rating classifications at 30 m versus 60 m showed a significant difference in the classification accu-racy for three dates (Fig. 2). These three dates were observed in 2018, where flights at 30 m had a better accuracy of classification. The remaining five dates across 2018 and 2019 did not have any difference in accuracy between the two flown heights (Fig. 2). The IDC classification accuracy ranged from 0.44-0.91. We noted that the first two dates in 2018 had lower accuracy than the remaining dates ranging from 0.44-0.64 whereas, the remaining time points had accuracy ranges from 0.71-0.91 (Fig. 2), which were similar to or better than previously reported (0.66-0.77) (Dobbels and Lorenz 2019).

**Figure 2:**
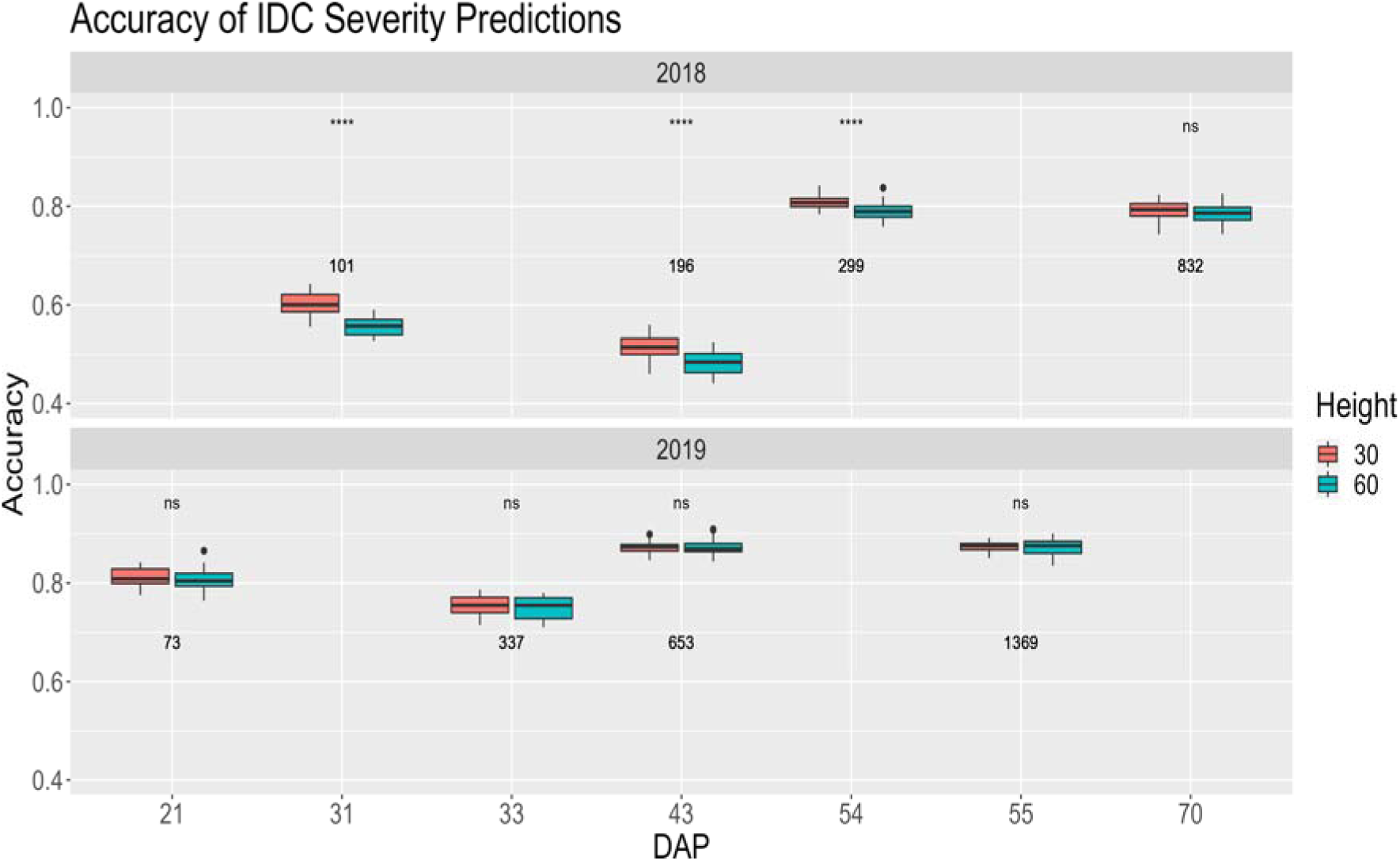
Comparison of the IDC classification accuracy between 30 m and 60 m flights.‘ns’ stands for not significant, and **** represents a p-value of ≤ 0.0001. The x-axis is the days after planting (DAP), and the y-axis is the accuracy of each IDC classification prediction. The top panel is 2018 and the lower panel is 2019. The numbers within the graph are the average canopy area for each flight date.

### 3.2 Effect of IDC on canopy area and growth

The time series data was primarily utilized to make two types of comparisons from a diverse collection of soybean accessions: (a) comparison of canopy areas per accession in the IDC field with the adjoining non-IDC field (Fig. 3a), and (b) 2018 IDC field observations versus 2019 IDC field observations (Fig. 3b). In Fig. 3, the x-axis is presented in DAP due to the differences in planting dates of the two fields, and the y-axis is the log transformation of the canopy area in meters^2^. We observed that the canopy areas were consistently higher in non-IDC field than in the IDC field, and was noted at each timepoint. The intercept varied by 0.76 (Fig. 3a). The difference in intercept shows that IDC symptoms start causing a reduction in the canopy area from an early stage, before the first flight measurements were made. Using observations of canopy area of accessions from the non-IDC field and IDC field, IDC symptoms show a 15.8% reduction in canopy area growth as observed by comparing the slopes. The change in slope signifies that the canopy area of accessions continues to be constrained in the IDC field up to 70 DAP, where we stopped taking further measurements. Consequently, we did not extrapolate past 70 DAP. When comparing canopy area results of 2018 and 2019, a linear growth pattern was observed with slopes of 0.053 for 2018, and 0.085 in 2019. In 2019, a steeper slope was observed (Fig. 3b), but it also experienced less severe IDC symptoms, which may have contributed to the larger growth rate in 2019. It is likely that the lower canopy area in 2018 is attributable to higher rainfall and associated water logging in the early season of that year. The accumulated rainfall from May 1*st* to August 31*st* was 689 and 446 mm in 2018 and 2019, respectively. In the first 30 days after planting, the rainfall was 290 mm in 2018 and 170 mm in 2019, signifying the excessive rainfall in the early season.

**Figure 3:**
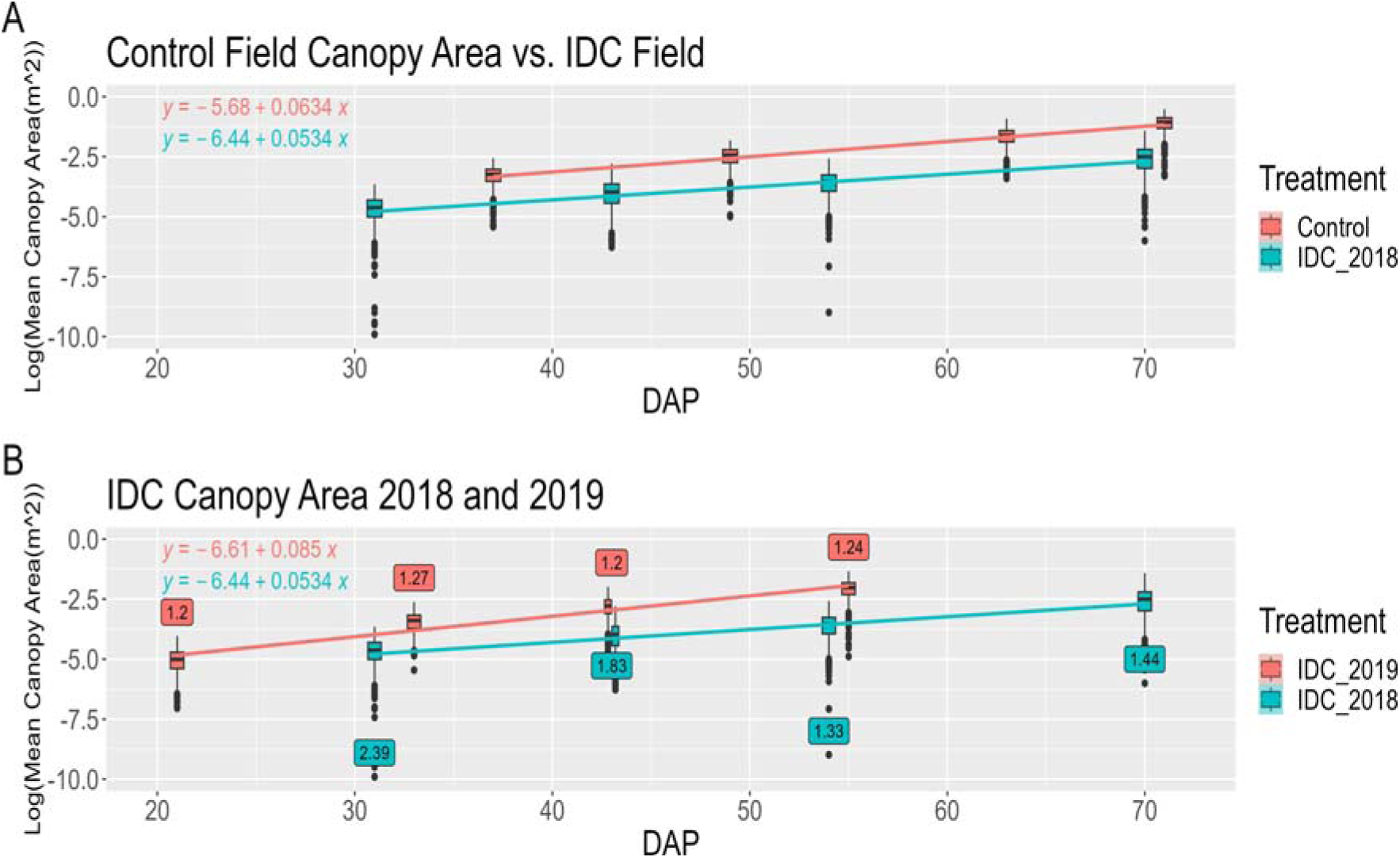
Linear regressions were calculated for IDC and non-IDC fields to estimate growth rates across timepoints. The x-axis is days after planting (DAP) for each time point and the y-axis is the log of the Mean canopy area for 477 diverse soybean accessions. (a) IDC and non-IDC field are compared in 2018. (b) IDC field in 2019 and 2018. Average IDC symptoms at each time point in year 2018 and 2019 in IDC fields, are shown in the box for each timepoint.

The effect of IDC on canopy area was also captured by comparing the near-isogenic pair of Clark (IDC tolerant) and IsoClark (IDC susceptible) (Fig. 4a and b). Since Clark and IsoClark are near iso-genic lines which have historically been used as checks for IDC tests in soybean, they are useful in highlighting the effect of IDC severity symptoms without the large genetic variance that we have across the rest of our panel (several hundred diverse accessions). Clark consistently had lower IDC scores compared to IsoClark (Fig. 4a) with an average IDC rating across all timepoints of 1.3 and 3.1, respectively. The IDC symptoms in Clark showed a rapid improvement (i.e., reduction in symptoms) in 2018. In 2019, Clark did not express any symptoms at any time points (Fig. 4a). In 2018 and 2019, Clark had a larger canopy growth rate than IsoClark (Fig. 4b). In both 2018 and 2019, IsoClark expressed symptoms at the first time point and did not recover in later time points, i.e., it maintained or increased IDC symptoms and showed reduced canopy area and canopy growth rate (Fig. 4b). The canopy growth rate for IsoClark was 32.5% and 46.7% less than Clark in 2018 and 2019, respectively.

**Figure 4:**
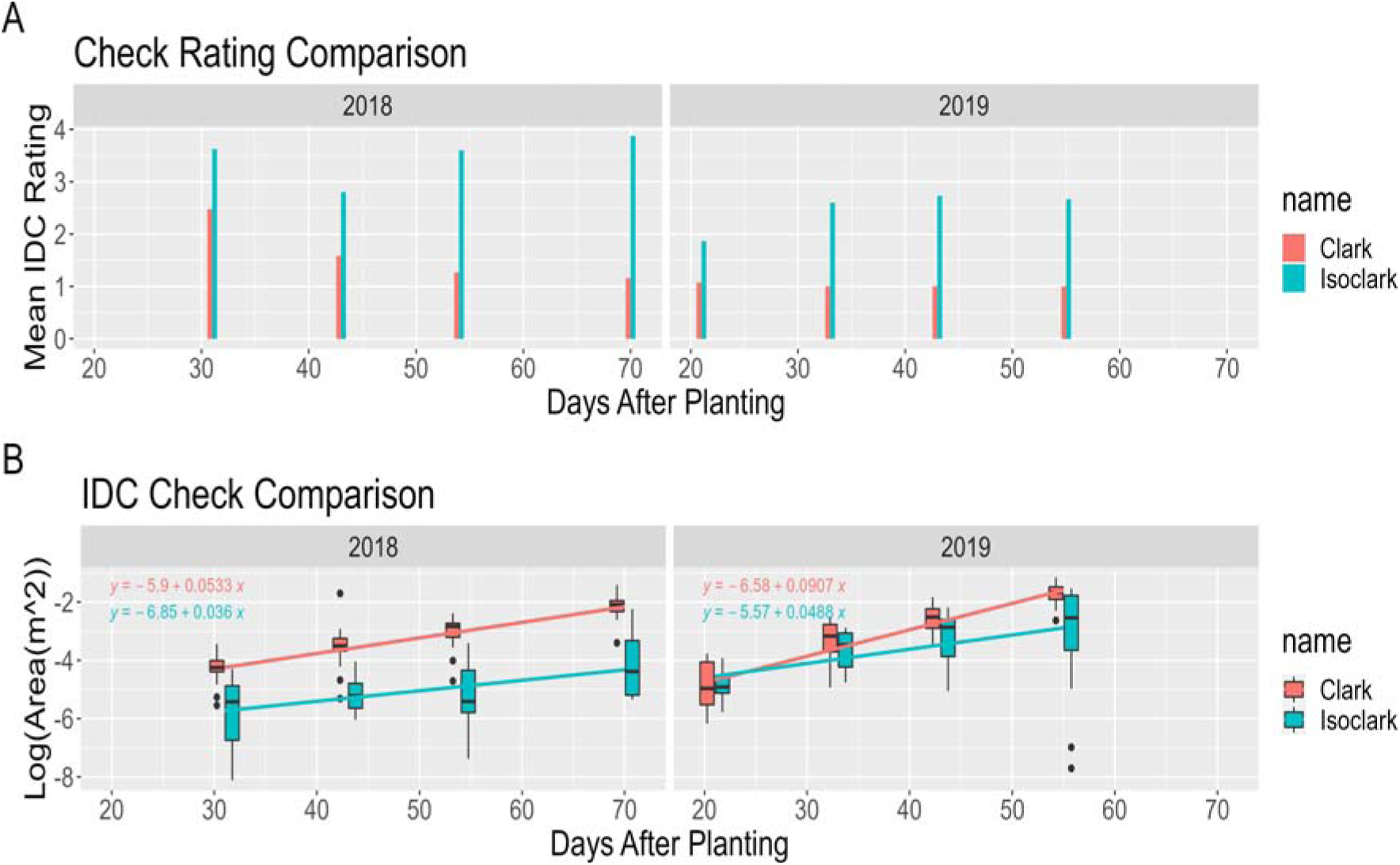
(a) IDC severity ratings across timepoints in Clark (IDC tolerant) and IsoClark (IDC susceptible) in 2018 and 2019. (b) Canopy area and canopy growth rate across time series in Clark and Iso-Clark.

IDC severity ratings from the IDC field and the baseline canopy area from non-IDC field had little to no correlation (maximum of 0.1), while a stronger negative correlation was observed among IDC severity ratings using 477 accessions and the canopy area of the same accessions from the IDC field (-0.4 to -0.3 across timepoints) (Fig. 5). Moderate correlation was noted for the canopy areas of the 477 accessions common across the IDC and non-IDC fields (0.2 - 0.3 across time points). Together these results indicate that IDC expression is independent of canopy area in non-IDC field; however, in fields with IDC suitable conditions, IDC ratings are inversely related to canopy area in the field (Fig. 5). Further credence was provided with the positive correlation of IDC ratings with a reduction in canopy area (from IDC compared to non-IDC field canopies), as shown in Fig. 5. Together, these results allude to an independent mechanism of canopy area development in IDC stressed and non-IDC stressed conditions.

**Figure 5:**
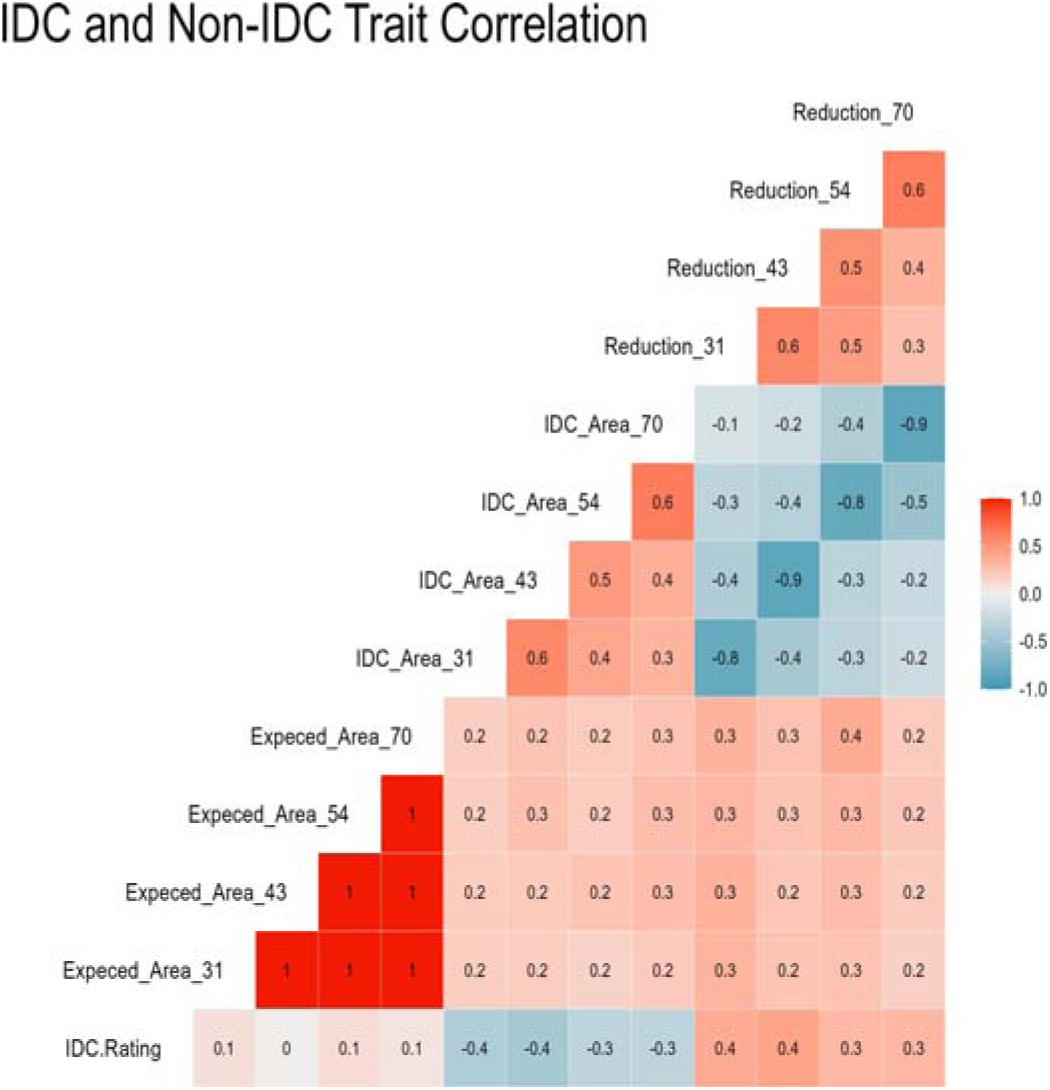
Trait correlation matrix for the control field (non-IDC field) and IDC field for IDC related traits. Data from four dates from 2018 IDC field and 2018 non-IDC field with 477 accessions were used to generate the correlation matrix. Key for traits: IDC Rating (IDC.Rating), Expected Canopy Area(Expected Area DAP #), IDC canopy area (IDC Area DAP #), and Canopy Area Reduction (Reduction DAP #).

The study of the relationship of IDC severity and canopy area each year showed a significant difference between IDC severity classes and the canopy area (Fig. 6). Phenotypic variance for canopy area was large at each IDC severity rating class. Across all timepoints the average canopy reduction was 9, 23, 29, and 58% when contrasting the average canopy area between each severity class. The loss in yield attributed to IDC has historically been estimated at 20% per change in rating classification. While canopy area and end season yield are not a 1:1 ratio, it appears that the loss of canopy area due to severity symptoms may follow a non-linear pattern with higher loss coming from larger severity ratings.

**Figure 6:**
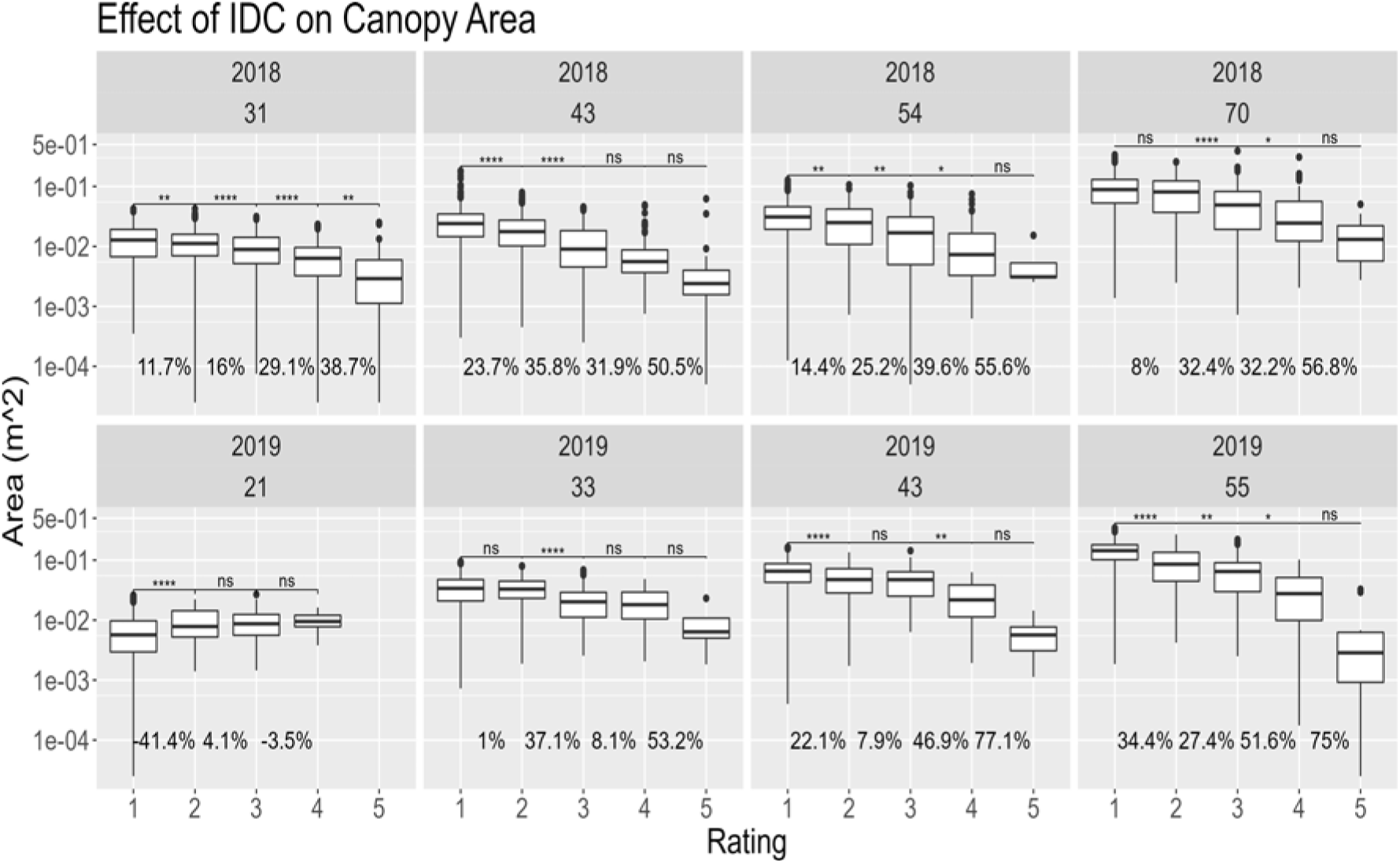
Pairwise comparisons of Canopy Area at each timepoint from the IDC field. In 2018, 608 diverse accessions were used for this study, and overlapping 580 soybean accessions were used in 2019. On the x-axis IDC severity rating is presented, and the y-axis is Canopy Area in m^2^. *, **,***,****, and ns represent p-values of ≤ 0.05, p ≤0.01, p≤ 0.001, p ≤ 0.0001, and p *>* 0.05. The value between each rating class is the average canopy reduction.

### 3.3 GWAS Results

GWAS identified 120 unique SNPs across the canopy and severity rating traits, which were significant after an FDR correction. Seventy-three unique SNPs were found for CGR, and 47 unique SNPs were found for IDC severity ratings across the manual, 30 m, and 60 m methods. Figure 7 shows the distribution of a subset of significant SNPs across timepoints for both traits, IDC severity and CGR.

**Figure 7:**
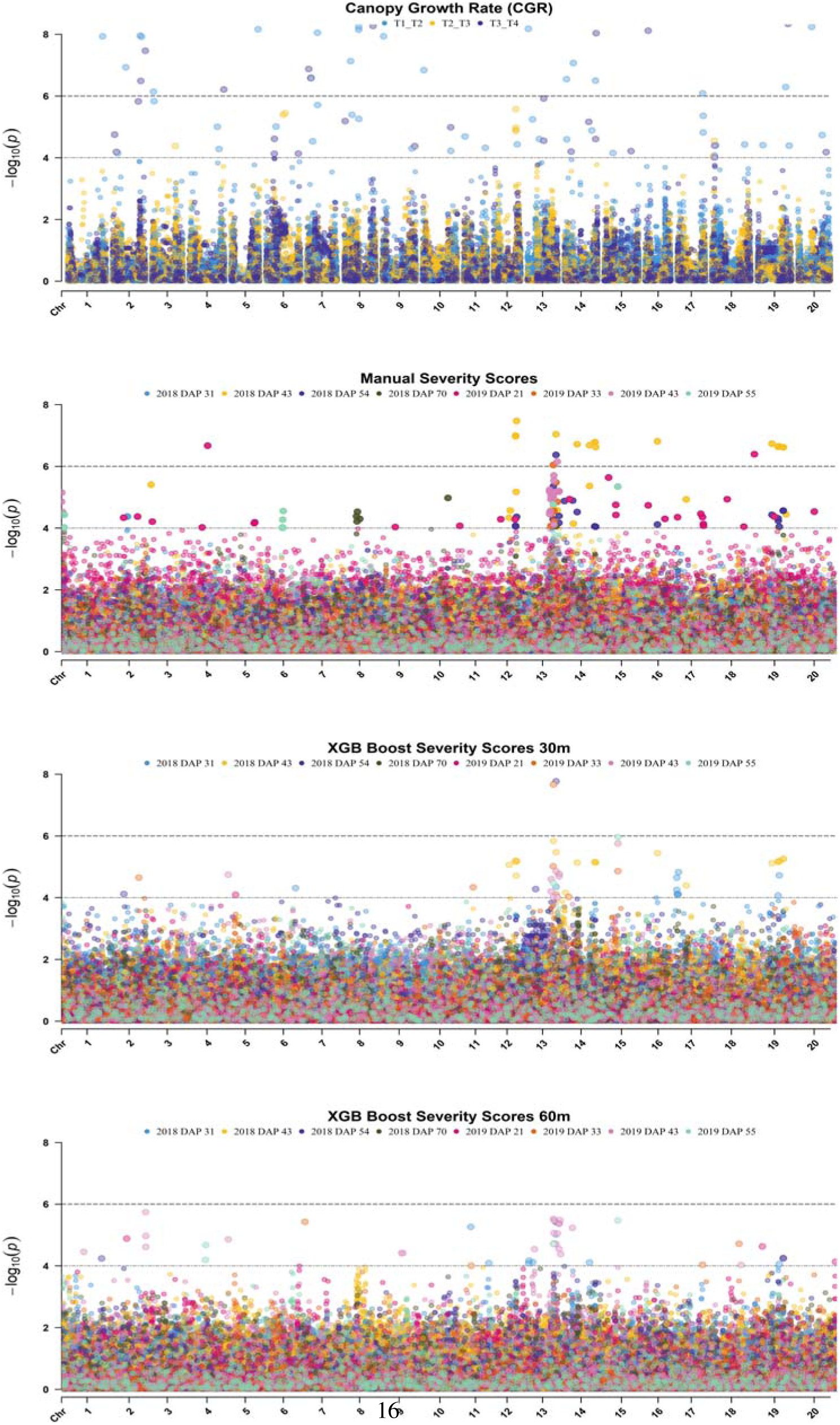
Significant SNPs identified and their distribution throughout the genome.

#### 3.3.1 IDC Severity

Across the three data collection methods (manual rating - control; 30 m and 60 m drone height images) 47 significant unique SNPs for IDC severity were detected. Over both years, the manual visual rating had the highest number of significant SNPs, and the 30 m height identified more significant SNPs than the 60 m flight (Table 1). The 30 m flight identified 20 significant SNPs in 2018, and only two significant SNPs in 2019. The 60 m flight identified no significant SNPs in 2018, and 13 SNPs were identified in 2019. The majority of the SNPs identified from the drone flights were also identified by the manual rating method. In 2018 for the 30 m flight, 18 of the 19 SNPs identified at 43 DAP were also detected with the manual method. For 54 DAP, only one SNP was identified via drone, and it was also identified via the manual ratings. No SNPs were identified for 31 or 70 DAP in 2018. In 2019, the one SNP identified by the 30 m flight at 33 DAP was also identified by the manual rating. This SNP is the same as the one identified by both manual and 30 m flights in 2018 at 43 DAP. An additional one SNP was detected for the 30 m flight at 55 DAP, which was not detected by the manual ratings, nor detected in 2018. The 60 m drone flight identified 13 significant SNPs at 43 DAP, of which seven were also detected by the manual ratings. The manual ratings were the only method that detected SNPs in 2019 at 21 DAPs. These results are summarized in Table 1.

**Table 1:**
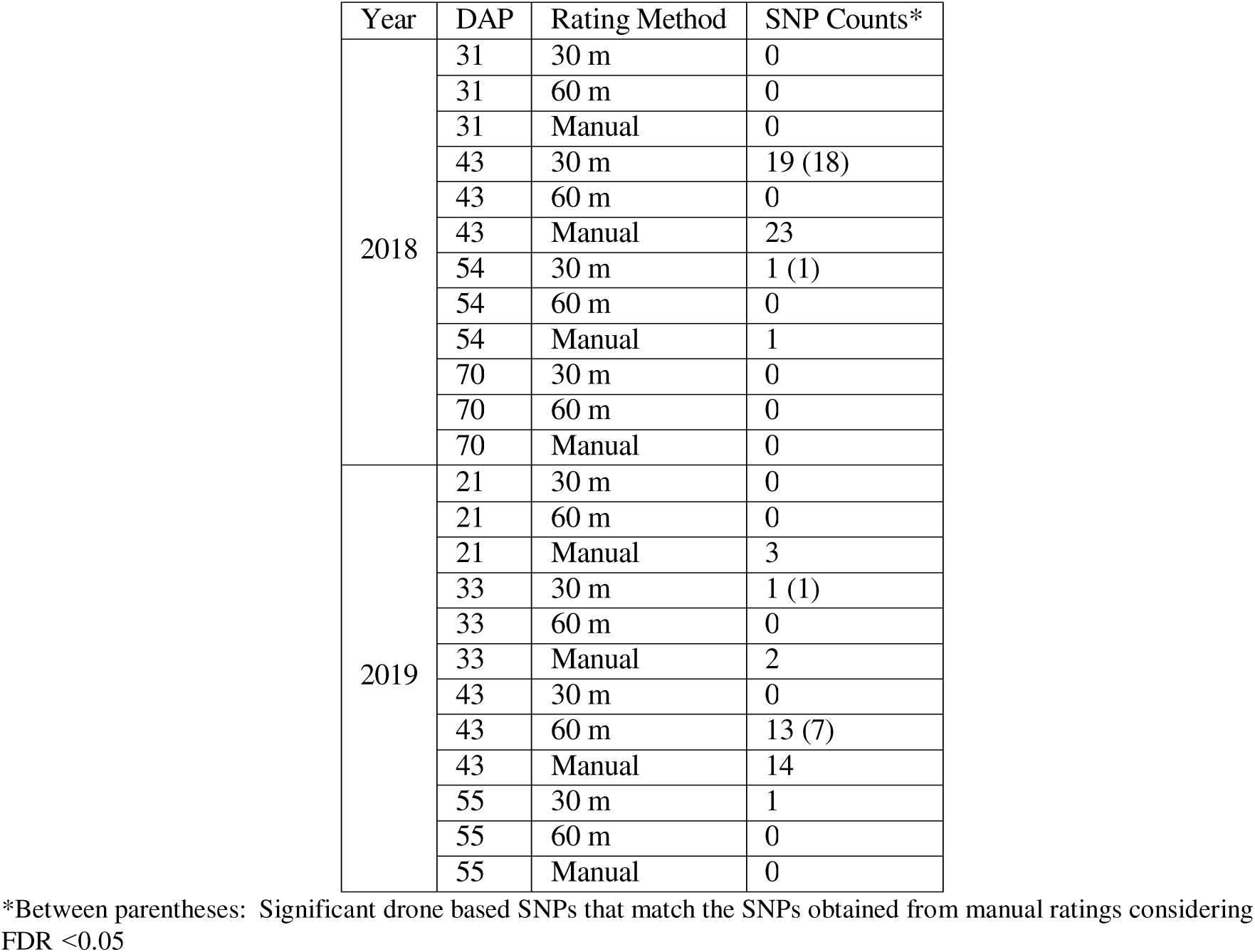
Summary of Significant SNPs found for each collection method of IDC Severity.

| Year | DAP | Rating Method | SNP Counts* |
| --- | --- | --- | --- |
| 2018 | 31 | 30 m | 0 |
|  | 31 | 60 m | 0 |
|  | 31 | Manual | 0 |
|  | 43 | 30 m | 19 (18) |
|  | 43 | 60 m | 0 |
|  | 43 | Manual | 23 |
|  | 54 | 30 m | 1 (1) |
|  | 54 | 60 m | 0 |
|  | 54 | Manual | 1 |
|  | 70 | 30 m | 0 |
|  | 70 | 60 m | 0 |
|  | 70 | Manual | 0 |
| 2019 | 21 | 30 m | 0 |
|  | 21 | 60 m | 0 |
|  | 21 | Manual | 3 |
|  | 33 | 30 m | 1 (1) |
|  | 33 | 60 m | 0 |
|  | 33 | Manual | 2 |
|  | 43 | 30 m | 0 |
|  | 43 | 60 m | 13 (7) |
|  | 43 | Manual | 14 |
|  | 55 | 30 m | 1 |
|  | 55 | 60 m | 0 |
|  | 55 | Manual | 0 |

\*Between parentheses: Significant drone based SNPs that match the SNPs obtained from manual ratings considering FDR <0.05

Of the SNPs identified across methods in 2018 and 2019, 15 SNPs have previously been reported as significant for IDC severity in soybean (Assefa et al. 2020; Merry et al. 2019; Jiaoping Zhang, Naik, et al. 2017), and are all located on Chr03 and Chr05. Using SoyBase (Brown et al. 2021) gene search tools, we found that 40 of the 47 significant SNPs have at least one known gene related to iron response, and are associated with IDC. A further investigation found that 22 of these genes have been reported as differentially expressed in low Fe and/or bicarbonate treatments (Waters et al. 2018). One gene on Chr04, *Glyma.04G209800*, is linked to two of the identified SNPs, *ss715588569* and *ss715588571*. This gene codifies to a transporter of γ-Aminobutyric Acid. In rice, γ-Aminobutyric Acid has been shown to supress iron transportation between the roots and shoots (Zhu et al. 2020). Three of the identified SNPs (*ss715615827*, *ss715615830*, and *s715615838*) are associated with four genes on Chr13: *Glyma.13G267400*, *Glyma.13G267500*, *Glyma.13G267600*, and *Glyma.13G267700*. These genes are homologs of the WYRK70 transcription factor, which plays a role in plant senescence and defense signaling (Yu et al. 2016). Another three SNPs (*ss715624341*, *ss715624357*, and *ss715625339*) were found to be linked to five genes on Chr16: *Glyma.16G147600*, *Glyma.16G147700*, *Glyma.16G149200*, *Glyma.16G149300*, and *Glyma.16G068100*, which are related to cytochrome C or P450 subunit assembly. Finally, *ss715638825* was found to be linked to *Glyma.20G236200* on Chr20, which is

#### 3.3.2 Canopy Growth Rate

In 2018, there were 73 significant SNPs identified across all time points for CGR. In 2019, there were no SNPs identified between T2-T3 and T3-T4. The QQ-plot for the 2019 T1-T2 was irregular (not presented), consequently the resulting significant SNPs were not investigated further. Therefore, further reported CGR results only come from the 2018 data. For the T1-T2 timepoint, 50 significant SNPs were identified, and an additional 23 SNPs were uniquely identified for the T3-T4 timepoint. No significant SNPs were found for the T2-T3 timepoint.

To study the association of CGR SNPs with iron deficiency or stress response, we performed a candidate gene search related exclusively to IDC or iron related responses. A summary of significant SNPs from 2018 on CGR are reported in Table 2 and includes *ss715582695*, *ss715591942*, *ss715598819*, *ss715607132*, and *s715607134*. In 2018, *ss715582695* was reported as significant for T1-T2, and is 3.4 kbp from *Glyma.02G222300*, which is a known NAC transcription factor that is upregulated in iron deficient conditions (O’Rourke et al. 2021). The SNP *ss715598819* is 73.56 kbp from the peak SNP *ss715598800* of a QTL identified in association with IDC (Assefa et al. 2020). *Glyma.07G067700* and *Glyma.16G033800* are both ferric reduction oxidase 2 genes (FRO2), and are near significant SNPs we detected. *Glyma.07G067700* is 43.6 kbp from the SNP *ss715598435*, while *Glyma.16G033800* is 19.0 kbp from the significant SNP *ss715624477*. FRO2 is a protein that has been found to be essential in iron homeostasis, and FRO2 genes have been previously identified in IDC studies (Mamidi et al. 2014; Moran Lauter et al. 2014). Across the two time points, T1-T2 and T3-T4, two significant SNPs, *ss715607134* and *ss715607132*, were identified that are 10.9 kbp apart from each other and near *Glyma.10G187000*, which has been found previously to be correlated with the expression of long-chain acyl-CoA synthases associated with stress response (J. Wang et al. 2022).

**Table 2:** SNPs associated with Canopy Growth Rate.

| Trait | SNP | Chromosome | Position | Region of interest | Annotation |
| --- | --- | --- | --- | --- | --- |
| 2018 T1_T2 | *ss715582695* | 2 | 40998890 | *Glyma.02G222300* | NAC transcription factor |
| 2018 T1_T2 | *ss715591942* | 5 | 39928655 | *Glyma.05G219400* | Heat shock protein |
| 2018 T1_T2 | *ss715598819* | 7 | 8414257 | 73.56 kbp from *ss715598800* | QTL peak associated with IDC |
| 2018 T1_T2 | *ss715607134* | 10 | 41985166 | *Glyma.10G187000* | Ethylene-responsive factor gene |
| 2018 T3_T4 | *ss715607132* | 10 | 41974315 | *Glyma.10G187000* | Ethylene-responsive factor gene |

## 4 Discussion

We studied IDC and canopy growth rate traits in soybean using drone based imaging at two different altitudes. This allowed us to compare the effect of flight altitude on the accuracy of IDC severity classification, as well as to study the effect the IDC severity ratings have on canopy size and growth during a portion of the growing season. For this, we built a customizable pipeline for using digital imaging techniques to concurrently study multiple traits, which we further applied by conducting a GWAS on the multiple timepoints of IDC severity and CGR to determine if temporal variations in significant loci existed.

### 4.1 Drone based imaging and data extraction tools

The drone based analysis pipeline presented offers a customizable solution to extract plot level data from orthomosaic images. The main difference that come from this pipeline compared to previous work is that it is based on the Esri (Redlands, CA) arcpy python library to run the analysis. The development of the plot files are also not dependent on the user rotating the image to draw a field bounding box that then generates the files for each plot (Matias et al. 2020). The difficulty in canopy area extraction in plant stress phenotyping is that the canopy pixels are not all as green as would be expected in non-stressed conditions, and a simple hresholding method is not applicable for canopy trait extraction. In our approach, we demonstrated the use of an unsupervised ISO-clustering classification method, which is similar to the K-means method that was used in previous work focused on IDC UAV classification (Dobbels and Lorenz 2019). The pipeline presented can be run using jupyter notebooks running python 3 and the method proposed applies to any field planted with a row-column design, and allows any plant scientists with stitched images to extract plot level data for downstream analysis. In comparison to FIELDImageR, a popular R-based tool for orthomosaic analysis for field trials, our plot extraction tool is based on python programming and does not have RAM memory limitations when running analysis, which can be an issue with large file sizes.

### 4.2 Accuracy comparisons for IDC severity classifications at 30m and 60m drone heights

When comparing accuracy of classification at 30 and 60m, there was a difference in the accuracy of classification for the first three timepoints in 2018. The average canopy area was 101, 196, and 299 cm^2^, respectively. There was no difference in the accuracy at the different heights in all other timepoints. While at he first timepoint of 2019, there was an average canopy area of 73 cm ^2^, all other timepoints had larger canopy area. This suggests that for canopy classification applications, there is a minimum threshold where higher resolution images are beneficial. This builds on the recommendations from Dobbels et al. on investigating the quality of classification at multiple flight altitudes, and is important in the implementation of drone based plant stress phenotyping at varying field size scales. The GSD at 30m is half the size of the GSD at 60m, and our hypothesis was that the accuracy would be greater at 30m due to the increase in resolution. If the quality of the image-based scores is equivalent at these heights, then the same sized field can be flown with the same flight parameters at 4x the speed. This means that a field that takes 60 minutes to fly at 30m will take approximately 15 minutes to fly at 60m. This also reduces the data size and processing time, because of the reduction in the number of pixels per unit area. This finding may apply to other plant stress phenotyping nurseries, but breeders should be mindful of what they need to detect to classify different stresses. If the stress is present on the whole canopy level then these findings should be transferable. However, if the stress signature is smaller, then lower flight altitudes may still be needed. Plant scientist should be sure to thoroughly examine the traits of interest before starting a new experiment to determine what parameters for data collection are needed to ensure that high quality and relevant data is being collected (Guo et al. 2021).

### 4.3 Effect of canopy size and growth on IDC ratings

Researchers have noted that plants are able to outgrow phenotypically visible IDC symptoms over time; however, our results indicates a reduction in canopy area even if the symptoms are no longer visible. Canopy size has been used in progeny row testing in soybean, and has been shown to be an effective selection tool (Moreira et al. 2019). Canopy area is an important physiological component trait for yield and the reduction in canopy size is a known effect of IDC stress in soybean. Reduction in leaf area as well as reduction in functional leaf area will contribute to a reduction in yield. This is done by reducing the plants ability to assimilate carbon to the plant through the photosynthetic process. This reduction has a direct impact on end season yield and the yield potential of a plant, especially in the reproductive phase (Vogel et al. 2021). With traditional phenotyping methods, a single discrete score is given to each plot, and this is used to assess severity. Raters can see variability in canopy area within rating classes, but this trait is not quantifiable without digital imagery. If a breeder is assessing data from a stress screening nursery using canopy size as a covariate for selection this may give a more informative decision-making tool than only using the severity ratings. This analysis could be taken a step further in future work and look at the ideal time to phenotype for canopy based stress symptoms and its effect on end season yield in soybean by looking at the correlation of stress ratings at multiple time points have, and could also be used to quantify the relationship between reduction in canopy area and yield reduction (X. Li et al. 2018). These findings agree with previous IDC work that has shown that there is a significant difference in yield for each change in severity rating (Fehr 1982), and with more recent work that has shown significant differences in canopy area in IDC severity ratings using UAVs (Dobbels and Lorenz 2019).

### 4.4 Time series GWAS and significant loci

We conducted a time series GWAS comparing the IDC severity scores and canopy growth rates across four timepoints in each year. The need for time series GWAS and genetic mapping studies helps to better understand the causal agents for plant growth and development through a growing season. Many studies have shown that different regions of the genome control for traits across the season. This type of analysis helps to identify regions of interest that can be manipulated for introgression into breeding program pipelines. A ime series GWAS can be conducted using manually collected traits, however most breeding programs do not have the resources to intensively measure necessary traits multiple timepoints throughout the season. With the use of HTP technologies, such as drones, the labor and time required to collect traits at a high temporal resolution is more feasible, and comes with the additional benefit that secondary traits such as canopy area can be collected simultaneously. We note that canopy area and reduction within a stressed field is a polygenic trait with no SNP explaining a major portion of the variation. We found a high degree of similarity between manually collected severity ratings, and drone imagery based severity ratings, with several of these SNPs being associated with previously reported IDC SNPs. In addition, one interesting finding from this work is that in an IDC stressed field the control for canopy growth and development seems to be independent of most reported known IDC related loci (Fig. **??**).

## 5 Conclusion and Future work

This research showed minimal differences in the accuracy of classifying IDC severities at two UAS flown heights, and 60 m flight had similar accuracy as 30 m flights while doubling the GSD. We presented the benefits and drawbacks of flying at different heights for plant IDC phenotyping. These results will be useful for scientists who are developing protocols for digital plant stress phenotyping when moving from ground based to aerial based phenotyping (Rairdin et al. 2022). We also presented a method for field plot boundary delineation, and explored the genetic control of multiple canopy related traits under iron deficient conditions. We report the usefulness of UAS based digital phenotyping for the concurrent estimation of plant stress and canopy growth. With the use of GWAS, we report that IDC stress expression seems independent of canopy growth development in a non-IDC stress field, which indicates that the IDC expression and canopy development are likely independently controlled. Our analysis and data extraction techniques focused on the use of 2-dimensional traits such as canopy area, but future work should investigate the use of 3-dimensional raits such as canopy volume (Chiranjeevi, T. Young, et al. 2021; T. J. Young et al. 2023), and the root-shoot interactions that are present between IDC severity and iron efficiency in plants. Previously, researchers have studied root traits in non-stressed conditions (Carley et al. 2023; Falk, Talukder Z Jubery, et al. 2020; Falk, Talukder Zaki Jubery, et al. 2020; Talukder Zaki Jubery et al. 2021); however, it is important to include root phenotyping in plant stress studies. Increased phenotyping capabilities will give researchers a better model of plant stress interactions. IDC severity ratings were taken in a known environment with a single stress; and future work should investigate the ability of UAV’s to identify and classify multiple biotic and abiotic plant stresses in a single field (Ghosal, Blystone, et al. 2018). This work focused on a plant stress that can be detected in the visible light spectrum, but prior work has shown that disease can be detected using hyperspectral imaging (Nagasubramanian, S. Jones, Sarkar, et al. 2018). As HTP capabilities continue to increase, it is paramount that researchers determine how to train models with limited resources in available labeled datasets effectively (Kar, Nagasubramanian, Elango, Nair, et al. 2021; Nagasubramanian, T. Jubery, et al. 2021; Nagasubramanian, Asheesh Singh, et al. 2022). Previous work has shown that sensors have varying abilities to predict traits in plants, and provide prescriptive recommendations (Chiozza et al. 2021; K. A. Parmley et al. 2019). As researchers continue to look into plant phenotyping it is important that they think about not just HTP in season, but also utilizing historical weather and genotypic data to better inform models (J. Shook et al. 2021), as well as integrating soil features that play a role in the plant response (Carroll et al. 2024). It is imperative that as machine learning continues to advance in agriculture that is not only used as a research tool, but developed in a way that farmers can also utilize this technology to optimize their farming operations. It is essential that new technology is co-developed with social scientists to increase the speed and rate of adoption (Balabaygloo et al. 2023). As data, AI and information is more readily available integrated models that optimize decision tools that are developed for farmers can help to implement management practices that increase profitability while reducing input costs (Sarkar et al. 2023).

## Abbreviations

The following abbreviations are used in this manuscript

DAP: Days After Planting
DL: Deep Learning
GSD: Ground Sampling Distance
GWAS: Genome Wide Association Study
HTP: High Throughput Phenotyping
IDC: Iron Deficiency Chlorosis
ML: Machine Learning
XGBoost: Extreme Gradient Boosting

## Acknowledgments

**General:** We thank staff and student members of SinghSoybean group at ISU, particularly Brian Scott, Will Doepke, Jennifer Hicks for their assistance with field experiments and phenotyping.

## Author contributions

A.S., A.K.S., M.E.C., B.G. and S.S. conceived the research; M.E.C. A.S. and A.K.S. performed experiments and data collection; M.E.C. and S.H. built machine learning models; M.E.C. nterpreted the results with inputs from A.R., L.V.L., A.F., A.S. S.S., B.G. and A.K.S.; A.R., A.F. and M.E.C. performed the GWAS analysis; M.E.C., L.V.L. and A.F. interpreted GWAS results with inputs from A.S. and A.K.S.; M.E.C. prepared the first draft with A.K.S. All authors contributed in the development of the manuscript, and reviewed the manuscript.

## Data Availability

Data and Codes will be made available publicly after acceptance of the paper through corresponding authors’ GitHub pages.

